# Crowding tunes 3D collagen fibrils and reveals matrix regulation of cancer cell morphogenesis

**DOI:** 10.1101/274696

**Authors:** A. Han, S. Ranamukhaarachchi, D. O. Velez, A. Kumar, A. J. Engler, S. I. Fraley

## Abstract

It is well established that the collagenous extracellular matrix surrounding solid tumors significantly influences the dissemination of cancer cells. However, the underlying mechanisms remain poorly understood, in part because of a lack of methods to modulate collagen fibril topology in the presence of embedded cells. In this work, we develop a technique to tune the fibril architecture of cell-laden 3D collagen matrices using PEG as an inert molecular crowding agent. With this approach, we demonstrate that fibril length and pore size can be modulated independently of bulk collagen density and stiffness. Using live cell imaging and quantitative analysis, we show that matrices with long fibrils induce cell elongation and single cell migration, while shorter fibrils induce cell rounding, collective migration, and morphogenesis. We conclude that fibril architecture is an independent regulator of cancer cell phenotype and that cell shape and invasion strategy are functions of collagen fibril length.

## Introduction

Accumulating evidence suggests that matrix architecture is capable of modulating cell migration phenotype ^1-4^ as profoundly as matrix stiffness ^5-7^. Largely, studies of matrix architecture have relied on micropatterned 2D surfaces and have focused on imparting contact guidance. Systematically controlling 3D ECM architecture remains a substantial challenge. Yet, it is now widely appreciated that cell behavior is distinct in native 3D ECM. Compared to 2D models, changes in the abundance, localization, and functional status of intracellular proteins have been documented in 3D culture ^8-10^. Thus, a major tradeoff exists between the physiological relevance of an ECM model system and the ability to tune and control specific physical features. It is currently impossible to decouple all of the architectural features of a 3D fibrilar protein network. Nonetheless, several studies have developed novel methods to produce highly aligned, anisotropic 3D collagen matrices, which impart both contact guidance and stiffness anisotropy ^11-14^. These methods include magnetic, mechanical, and cell force driven reorganization of collagen fibrils as well as electrospinning. From these studies, matrix stiffness and alignment have been established as modulators of cell phenotype through mechanotransduction processes ^15^. However, the mechanisms by which matrix architecture may act independently on cells are not clear. This is due in part to the scarcity of *in vitro* experimental techniques that satisfy the need to modulate fibril characteristics independently of collagen density and stiffness while also allowing cells to be fully embedded in 3D.

Molecular crowding (MC) is one approach that can potentially achieve these goals. Crowding is a physiologically relevant phenomenon, whereby high concentrations of macromolecules occupy the extracellular space and generate excluded volume effects^16,17^. In the context of collagen polymerization, this results in alterations to the rates of nucleation and fibril growth^18^.

Thus far, architectural engineering of 3D collagen hydrogels with MC has been investigated in cell-free conditions. Here, we establish a crowding technique for cell-laden 3D collagen matrices using biologically inert PEG. We demonstrate that adjustments to the amount of PEG added during collagen assembly and cell embedding reliably tune fibril topography. We then quantitatively evaluate the biophysical properties of the MC matrices as well as the morphological and migration response of cancer cells in the engineered constructs. Importantly, we show through control experiments that the influence of the crowding agent on cell morphology and migration behavior occurs through the topographic alterations MC induces in the matrix. Finally, we demonstrate that matrix architecture is a critical modulator of cancer cell phenotype independently of matrix stiffness or density.

## Results

### Macromolecular crowding with PEG tunes collagen fibril characteristics

To explore the impact of fibril architecture on cancer cell behavior in a 3D collagen matrix, we sought to develop a method for tuning fibril length in a collagen I hydrogel while simultaneously embedding cells. More specifically, our goal was to shorten fibril length of a 2.5 mg/ml collagen matrix without changing the density or stiffness of the matrix. The assembly of collagen I solution *in vitro* into a fibrous 3D matrix is thought to be driven by diffusion-limited growth of nucleated monomers (Fig. 1A)^19,20^. Crowding during collagen polymerization by the addition of 25 mg/ml of 400 Da Ficoll has previously been shown to tune fibril growth rate and architecture in cell-free conditions^18^. However, studies suggest that Ficoll is cytotoxic^21^. Thus, we sought to test PEG as a MC agent for its biological inertness.

**Figure 1.**
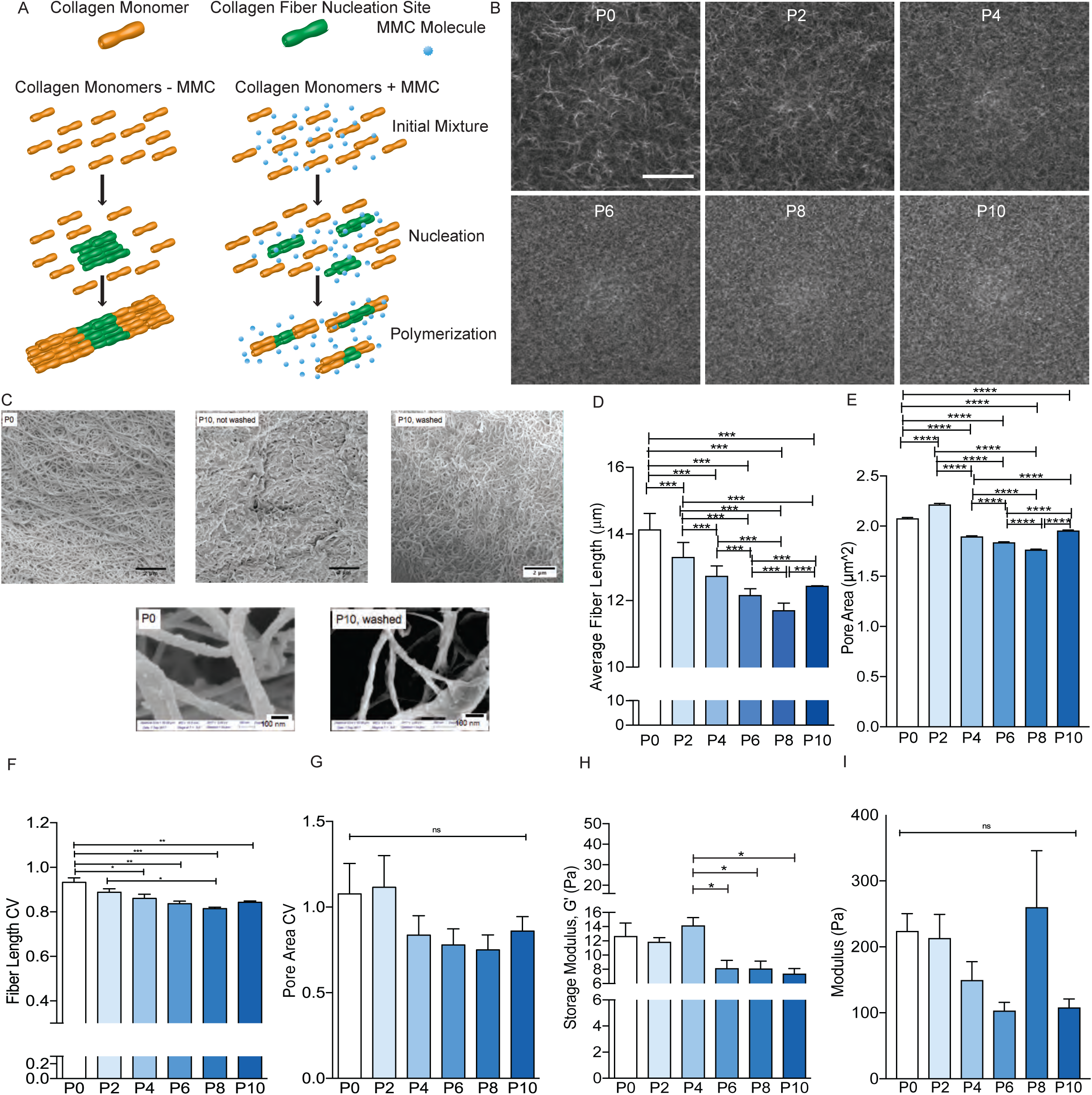
Fibril topography modulation by molecular crowding. (a) Schematic showing how molecular crowding affects matrix polymerization. (b) Reflection confocal micrographs of 2.5 mg/ml collagen polymerized without a molecular crowding agent, P0, or with 2-10 mg/ml of 8kDa PEG as a crowding agent, P2-P10. Scale bar is 200 μm. (c) SEM images of a 2.5 mg/mL collagen gel (top left) and 2.5 mg/ml collagen gels polymerized with 10mg/mL PEG without washing (top middle) or with thorough washing before fixing (top right). Bottom images are magnified versions of top left and right images. (d) Characterization of mean fibril length and (e) pore size as a function of the extent of crowding. (f) Coefficient of variation of fibril length and (g) pore size as a function of the extent of crowding. (h) Bulk elastic moduli of control and crowded matrices measured by shear rheometry. (i) Local moduli of control and crowded matrices measured by AFM. N=3 replicates for each condition. At least three fields of view were analyzed per replicate. Bar graphs show the mean and standard error of measurements. Statistical significance tested by ANOVA and reported as p<0.001, ***; p<0.01, **; p<0.05, *.

First, we tested the ability of PEG to alter collagen architecture. To do so, we introduced increasing amounts of 8,000 Da PEG (0-10 mg/ml, labeled P0-P10) into a 2.5 mg/ml collagen I solution, then washed the gel after polymerization to remove the PEG (see Methods), and finally imaged the resulting fibril architecture with reflection confocal microscopy. Increasing amounts of PEG led to gradual changes in collagen fibrils (Fig. 1B). To ensure that our washing procedure effectively removed the PEG, we conducted SEM imaging (Fig. 1C) of the P10 matrix with and without washing prior to fixation and sample processing for SEM (see Methods). We also imaged the P0 matrix for comparison (Fig. 1C). These images show that PEG is not detectable in the collagen matrix after washing. Quantitative analysis of reflection confocal images of the crowded matrices revealed a linear decrease (r^2^=0.73, p=0.03) in average fibril length from 14.1μm without PEG to 11.7μm with 8 mg/ml of PEG mixed in during polymerization (Fig. 1D). Interestingly, this trend reversed between 8 and 10 mg/ml of PEG, where average fibril length increased slightly from 11.7 μm to 12.4μm (Fig. 1D). The average pore size of the crowded matrices changed only slightly across all conditions, ranging from 1.75 to 2.2 μm^2^ (Fig. 1E). Average fibril width, analyzed by SEM, varied marginally (∼35 nm) with PEG crowding (Supplementary Fig. 1, A and B). These analyses indicated that although multiple matrix characteristics change simultaneously as a result of crowding, each feature follows its own characteristic dose-response relationship.

We also sought to quantify the relative heterogeneity of the fiber architecture in each condition to determine if this relates to the heterogeneity of cellular behavior in these matrices. Using the coefficient of variation (CV) to assess heterogeneity in the fibril length and pore size of each condition, we found that increased crowding results in a gradual decrease in CV (Fig. 1, F and G). This indicates that in general, crowding homogenizes the matrix architecture. However, we do observe a slight increase in CV for the most crowded condition, 10 mg/ml PEG, as was observed for average pore size and fibril length.

To further characterize the biophysical properties of the crowded collagen constructs, we measured their bulk and local elastic moduli using shear rheology and atomic force microscopy (AFM), respectively. An AFM tip of approximately the same size scale as mechanosensory nascent adhesions was chosen for this measurement^22^. Slight differences in the bulk moduli were observed. These were statistically significant between the P4 crowded condition (14 Pa) and higher crowding conditions (∼8Pa, Fig. 1H). However, no significant differences in local moduli were observed when averaged over multiple locations and biological replicates (Fig. 1I). Thus, overall, matrix architecture was tuned without altering matrix stiffness and without changing the density of the collagen. This similarity in mechanical behavior may result from a balance between the increase in the connectivity of the network and a simultaneous weakening of the strength of the connections^23^. It is important to note that the stiffness of 2.5 mg/ml collagen (here P0) has previously been shown to mimic normal breast tissue^7^. All of our PEG crowded and non-crowded 2.5 mg/ml collagen constructs are within this range of stiffness and can be considered representative of the mechanical conditions cancer cells encounter during invasion and metastasis.

### PEG crowding alone does not directly influence cell morphology or migration behavior

Having confirmed that PEG is an effective MC agent for tuning collagen architecture, we next sought to determine whether it would directly influence cell behavior independently of its effects on matrix architecture. First, we verified that PEG functions as an inert crowding agent in our system. Using fluorescently labeled PEG and a bottom-reading microplate reader, we measured the diffusion of the crowding agent into collagen gels and the effectiveness of our washing of the matrix. Supplementary Figure 1C shows that the fluorescence intensity, detected from the bottom of the well containing the matrix, displayed a time-dependent increase as fluorescent PEG was added on top and diffused into the gel. Likewise, when we conducted washing steps to remove the PEG, the fluorescent intensity at the bottom of the gel decreased, nearing the original baseline reading. Next, we conducted an additional control experiment to ensure that even if our washing procedure did not remove all traces of PEG from the matrix (as suggested by the SEM images in Figure 1C and the fluorescent PEG readings in Supplementary Figure 1C) cell behavioral differences result from fibril changes not cell-PEG interactions. To test this, we embedded MDA-MB-231 breast cancer cells in a 2.5 mg/ml collagen matrix with no PEG added during polymerization. Then, we added 10 mg/ml PEG, the maximum amount of PEG used for our matrix engineering experiments, on top of the fully polymerized matrix and allowed the 8,000 Da molecules, radius of gyration ∼3 nm^24^, to freely diffuse into the interstitial spaces (Fig. 2A). In these experiments, no washing of the matrices was conducted. Reflection confocal analysis of the matrix architecture with and without PEG added on top revealed very slight differences in average fibril length (<1μm) and pore size (<0.5 μm^2^) (Supplementary Fig. 1, D and E). It is important to note that the large number of pores and fibrils analyzed tends to generate statistical significance between conditions, even when differences are small^25^. After 15 hours of culture in this control condition, no significant differences were observed in cell morphology or migration, as assessed by cell circularity and the total path length traveled by the cells over the first 15 hours respectively (Fig. 2, B and C).

**Figure 2.**
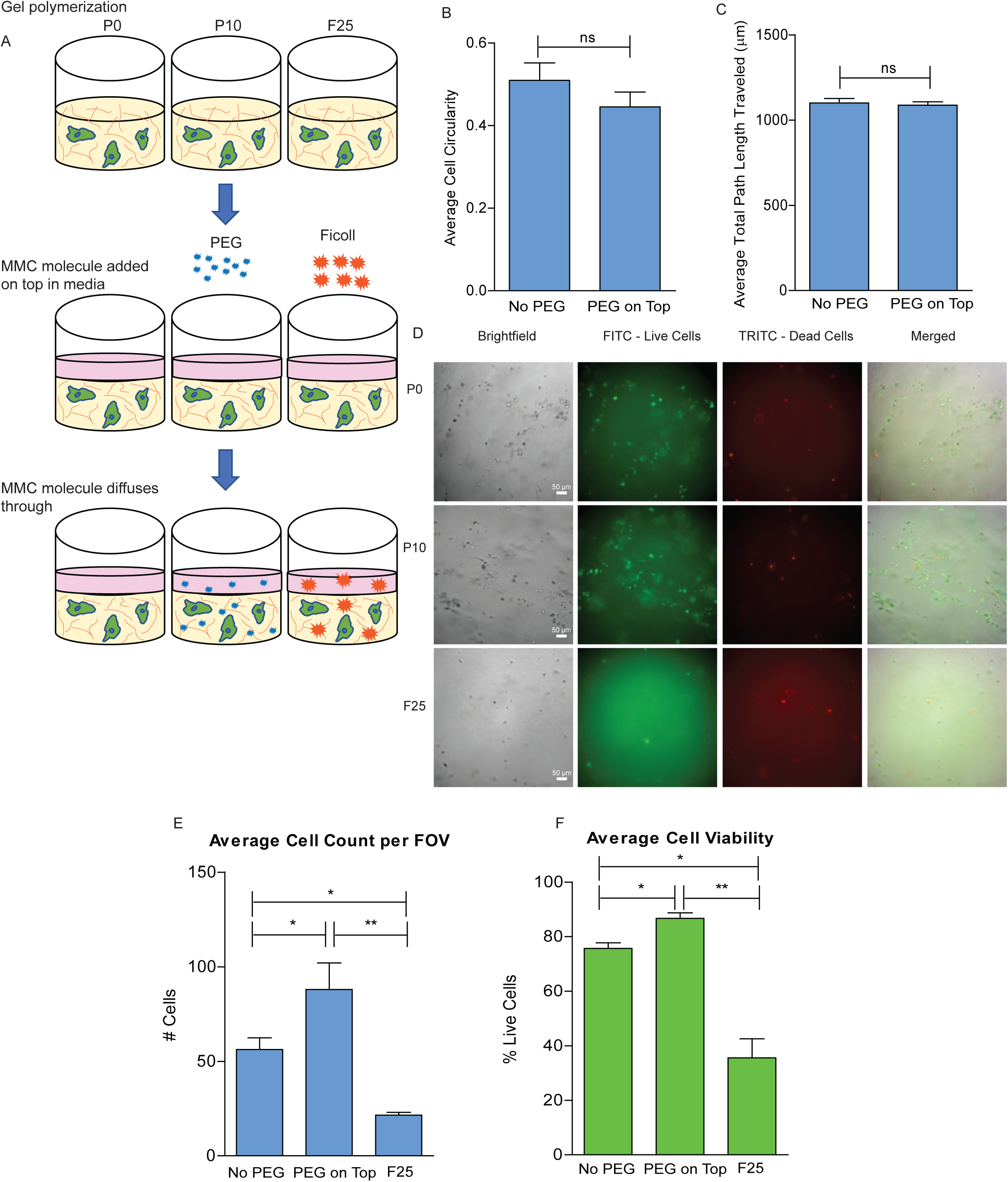
Influence of PEG crowding alone on cell morphology, migration, and viability in 3D. (a) Schematic of control experimental setup. PEG or Ficoll was added *after* collagen polymerization to evaluate potential effects on cell behavior independent of matrix changes. Influence of PEG crowding on (b) cell shape and (c) cell migration over 15 hrs. (d) Representative micrographs of cells after one week in culture showing brightfield (left), live (green) and dead (red) cell staining. Merged image on right. (e) Cell proliferation and (f) viability evaluated after one week of PEG or Ficoll crowding *after* polymerization. N=3 biological replicates for each condition. At least 100 cells were analyzed per condition. Bar graphs show the mean and standard error of measurements. Statistical significance tested by ANOVA and reported as p<0.001, ***; p<0.01, **; p<0.05, *.

Next, we assessed the viability of the cells after one week of culture in the control and crowded conditions where the MC agent was not washed out. For comparison, we also tested the effects of 25 mg/ml Ficoll 400 (400,000 Da) under the same condition, which has been used previously to tune collagen matrix architecture to approximately the same degree as we accomplish here using 10 mg/ml PEG^18^. Cells were seeded at the same initial density in all conditions. Figure 2D shows micrographs of cells after one week. Total cell count was significantly lower in the Ficoll crowded conditions compared to the non-crowded and PEG crowded conditions after one week, indicating that Ficoll negatively impacted cell proliferation while PEG did not (Fig. 2D, left column, and Fig.2E). Live-dead staining also revealed that cell viability was significantly reduced in Ficoll crowded conditions (Fig. 2F). Since Ficoll negatively impacted cell viability while PEG did not, and both achieved comparable changes to matrix architecture, we conclude that PEG crowding is a more useful technique to alter the fibril architecture for embedded cell studies.

### 3D collagen fibril topography patterns cell shape

Having established PEG crowding as a method to modulate collagen fibril topology, we next sought to quantify the influence of matrix architecture on the morphology and migration of embedded cancer cells. To do so, we polymerized 2.5 mg/ml collagen with a low seeding density of MDA-MB-231 cells and 0-10 mg/ml PEG mixed in. After polymerization, the cell-laden gels were washed to remove the PEG. Single cells were then monitored by timelapse microscopy. Figure 3A highlights typical cell morphology differences observed in the crowded matrices as representative cell outlines in each condition after 15 hours. These trends in cell shape were stable throughout the first 15 hours following matrix polymerization and washing (Supplementary Fig. 1F). Since cells were seeded sparsely and most had not yet divided during this time period (Supplementary Fig. 1G), these morphology differences result from cell-matrix interactions as opposed to cell-cell interactions. Quantitative assessment of individual cell shapes at 15 hours revealed that cell circularity follows a similar trend as fibril length. As fibrils were shortened by increased molecular crowding (Fig. 1D), cells became more rounded and less elongated (Fig. 3B) following a trend similar to that of fibril length. Mean, median, 75% values, and 25% values of fibril length each significantly predicted that of cell circularity (Pearson Correlation, Supplementary Fig. 2, A-D). As another test of this relationship, we would expect the CV of fibril length and the CV of cell circularity to follow a similar trend as well. That is, as fibril length becomes more homogenous with increased crowding, cell circularity would likewise become more homogenous. Indeed, as increased crowding causes lower CV of fibril length (Fig. 1F), cell circularity CV also decreased (Fig. 3C), although differences were not significant. The relationship between the CV of fibril length and CV of cell circularity was also significantly linear with a Pearson r=0.91 and p = 0.01 (Supplementary Fig. 2E). Next we compared the mean, median, 75% values, and 25% values of cell circularity to those of pore size and found that there were no significant relationships (Supplementary Fig. 3, A-D). These findings suggest that cell shape is a function of 3D fibril length.

**Figure 3.**
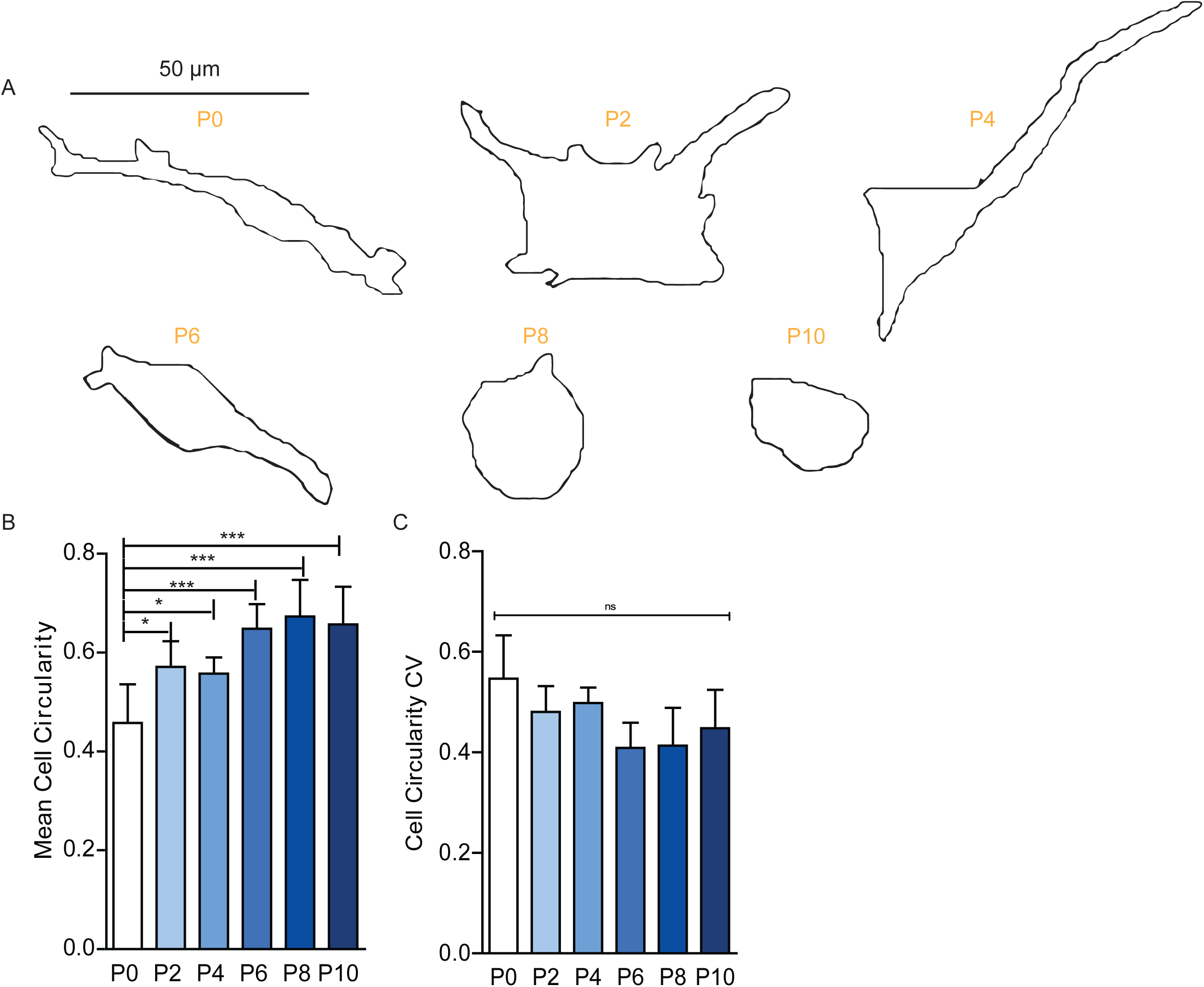
Influence of crowded collagen fibril architectures on cell shape in 3D. (a) Outlines of representative cells in each matrix condition, P0-P10, after 15 hours. (b) Mean cell circularity in each matrix construct. (c) Coefficient of variation of cell circularity in each matrix construct. N=3 biological replicates for each condition. At least 100 cells were analyzed per condition. Bar graphs show the mean and standard error. Statistical significance tested by ANOVA and reported as p<0.001, ***; p<0.01, **; p<0.05, *.

### 3D collagen fibril topography modulates a transition from single cell migration through collective cell migration to morphogenesis

To examine the impact of collagen fibril topography on breast cancer cell migration, we monitored MDA-MB-231 cells for one week in each construct. A striking transition from single cell migration to collective migration was observed in the 2.5 mg/ml collagen matrices crowded with 6 mg/ml of PEG (P6) and higher. Figure 4A shows representative micrographs of the breast cancer cells in each construct. Even more surprisingly, in P8 and P10 conditions the chain-like structures became more fused and smooth-edged, and other multicellular structures emerged at low frequency (Fig. 4, B and C). These structures resembled lobular and glandular structures of normal breast tissue^26^. Figure 4C shows the frequency of single cell, multicellular chain, and multicellular smooth structure phenotypes. It is important to reiterate that SEM imaging and fluorescent PEG experiments confirmed that our washing procedure effectively removed PEG after polymerization (Fig. 1C and Supplementary Fig. 1B), and further, the presence of PEG added on top of the matrix in control experiments did not impact cell behavior (Fig. 2, B and C). Thus, we can deduce that matrix architecture drives these phenotypic transitions, from single cell migration through collective migration to morphogenesis.

**Figure 4.**
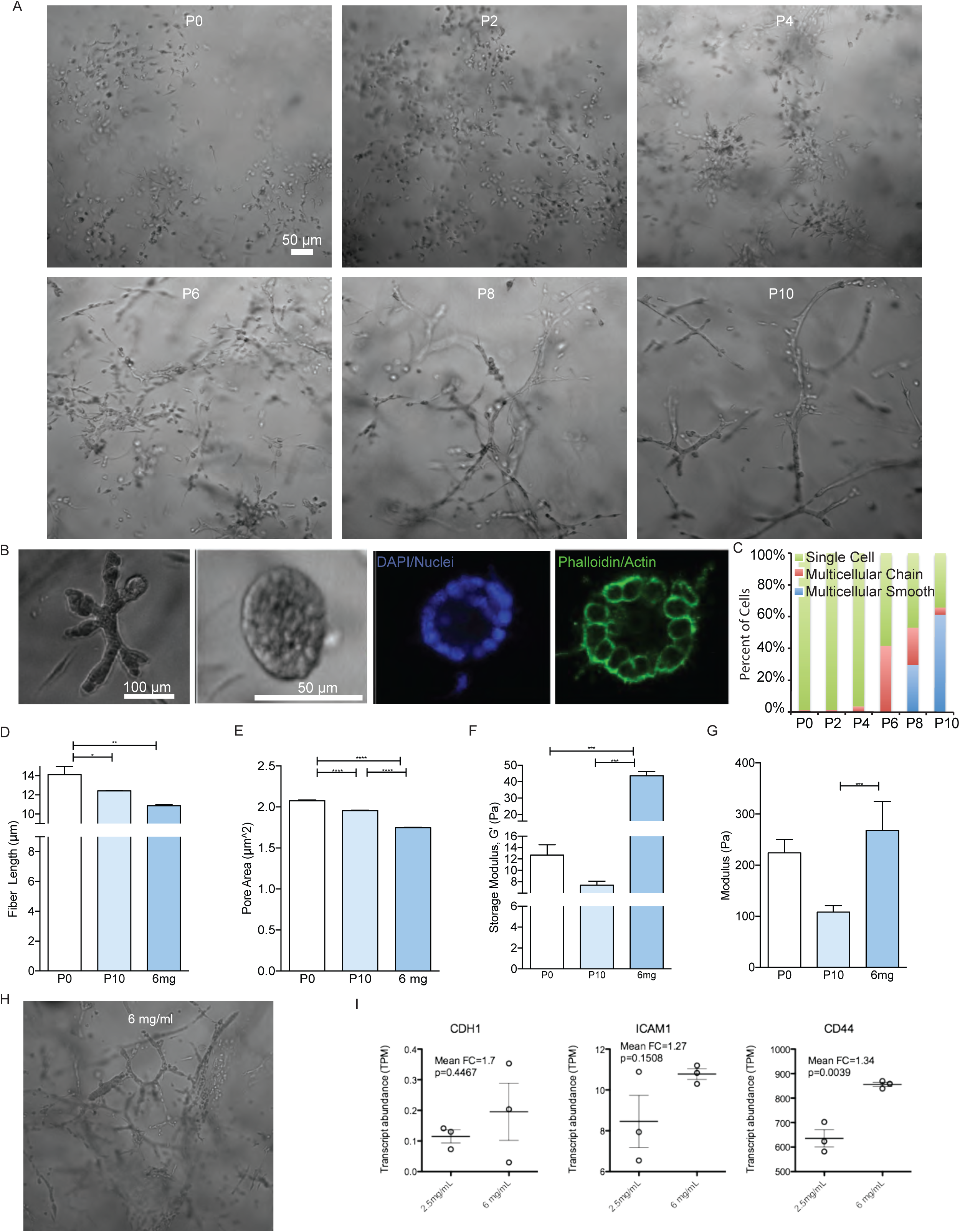
Influence of fibril topography on cell migration behavior in 3D. (a) Representative micrographs of cells in each matrix construct after one week. (b) Additional multicellular structures observed at low frequency in P8 and P10 conditions. Lobular (left) and acinar (right three images) structures resembling normal breast structures. Rightmost two images show representative acinar structure stained with DAPI (nuclei, blue) and phalloidin (actin, green) and reveal an organized and hollow morphology. (c) Frequency of phenotypes observed in each matrix construct. (d) Fibril length of high density 6 mg/ml collagen matrices compared to P0 and P10 conditions. (e) Pore size of 6 mg/ml collagen matrices compared to P0 and P10 conditions. (f) Bulk elastic modulus, measured by shear rheology, of 6 mg/ml collagen compared to P0 and P10 conditions. (g) Local modulus, measured by AFM, of 6 mg/ml collagen matrices compared to P0 and P10 conditions. (h) Micrographs of MDA-MB-231 cells forming network structures in high density 6 mg/ml collagen. (i) Transcript expression of cell-cell adhesion genes CDH1, ICAM1, and CD44.

To further confirm that PEG had no direct effect on cell behavior, we asked whether we could create a matrix of the same crowded architecture without using PEG. We hypothesized that a self-crowded (i.e. higher density) collagen solution would form a similar architecture to a PEG crowded collagen solution. Indeed, the architecture of a 6mg/ml collagen matrix resembled that of the P10 condition in terms of fiber length and pores size (Fig. 4, D and E). However, the bulk stiffness and local stiffness of the 6 mg/ml collagen matrix were significantly higher (Fig. 4, F and G). Nonetheless, in this architecture-matched matrix, MDA-MB-231 cells formed multicellular structures (Fig. 4H). Using RNA-seq analysis, we found that the multicellular phenotype was associated with upregulated expression of several cell-cell adhesion genes: CDH1, ICAM1, and CD44 (Fig. 4I).

We next sought to identify which matrix feature(s) could be responsible for driving the switch from single to collective migration and further morphogenesis in our PEG crowded constructs, where the total density and overall stiffness of collagen was held constant. Since the phenotypic transition of breast cancer cells in our constructs was not gradual but sharp, we hypothesized that matrix feature(s) could act in a thresholding capacity. To assess this, we compared characteristic values for each matrix feature to the frequency at which we observed single versus multicellular phenotypes across the matrix conditions. Plotting the mean, median, and CV of fibril length against the frequency of single cell migration revealed that indeed a threshold in fibril length predicted the emergence of multicellular phenotypes in P6-P10 (Fig. 5, A-C). Likewise, an associated cell circularity threshold was identified (Fig. 5D), reinforcing the relationship between fibril length and cell shape. However, pore size could not reliably threshold the phenotypic switch (Fig. 5, E and F), even when only the largest 50% of pores were considered (Fig. 5G). If only the maximum pore size was considered for each condition, a pore size threshold of ∼35 μm^2^ (or ∼7μm diameter) could be defined to separate single cell (P0-P4) from multicellular (P6-P10) conditions (Fig. 5H). However, these maximum pore sizes are not representative of the majority of the matrix architecture. Moreover, protrusions are often smaller than this threshold pore size (Fig. 3A), such that it would not limit cell spreading. Yet a significant decrease in cell spreading (increase in circularity) is observed at this threshold and is maintained for at least 15 hrs (Fig. 3B). Most likely, then, the correlation between cell circularity, fibril length, and collective morphogenesis represents a change in fibril adhesiveness or degradability.

**Figure 5.**
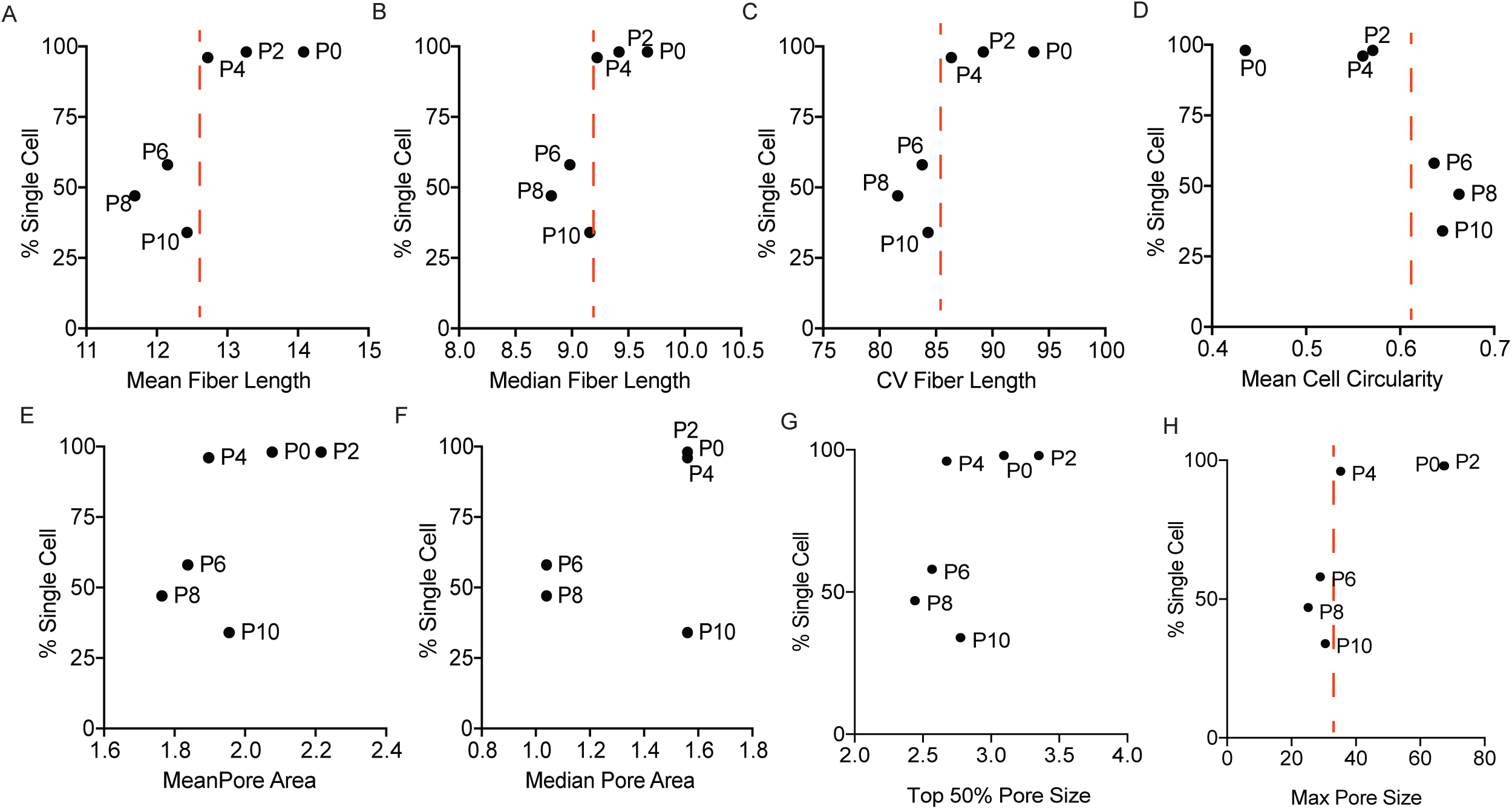
Matrix features predict cell shape and behavior. (a) Mean, (b) median, and (c) coefficient of variation of fibril length in each matrix construct plotted against the frequency of the single cell phenotype in each construct. Red dotted lines indicate fibril length threshold, below which cells transition into multicellular phenotypes. (d) Frequency of the single cell phenotype in each matrix construct plotted against the mean cell circularity in each construct. Red dotted line indicates threshold value below which cells transition into multicellular phenotypes. (e) Mean, (f) median, (g) top 50%, and (h) max pore area measurements plotted against the frequency of the single cell phenotype. N=3 biological replicates for each measurement. At least 300 cells were analyzed in each condition.

## Discussion

We created a novel 3D collagen system that physically decouples both stiffness and density from fibril architecture to independently assess the impact of fibril architecture on cell behavior. Our study reveals that when individual cells interact with different collagen matrix architectures, cell shape is a function of fibril length. Further, this interaction ultimately drives distinct modes of cell migration. Single cell migration is favored in matrices with long fibrils whereas multicellular cell migration and morphogenesis is favored in matrices with short fibrils. These two behaviors can be predicted based on a fibril length threshold and a related cell circularity threshold.

Previous work by Foty and Steinberg, using hanging drops and 2D systems, demonstrated that cell-ECM adhesion competes with cell-cell cohesion following physical principles related to surface tension^28^. It is possible that the short collagen fibrils in our system restrict the size or stability of cell-ECM adhesion compared to longer fibrils and thereby promote cell-cell cohesion. Alternatively, reduced degradability of the fibrils could be responsible for confining cells in a rounded shape. This could alter the tensegrity of the cell, reducing the activation of cell surface integrins and their affinity for binding ECM^29,30^. Tensile forces that are generated by contractile actomyosin filaments are resisted inside the cell by microtubules and outside the cell by the ECM and by adhesions to nearby cells^30^. In rounded single cells, microtubules serve as the primary resistance to the pre-stressed cytoskeleton and also provide a mechanical force balance to a tensed network of chromosomes and nuclear scaffolds^30^. This mechanical linkage could alter gene expression and cell behavior in a cell shape dependent way. Previous studies on 2D patterned substrates have shown that cellular geometry influences modular gene expression programs, differentiation, nuclear deformation, cytoskeleton reorganization, chromatin compaction, growth, apoptosis, and cell division ^31-36^. However, cell roundedness due to loss of attachment has also been shown to impair glucose uptake, inducing metabolic defects that drive gene expression changes^37^. It is also possible that our altered matrix enhances the sequestration of soluble factors and thereby autocrine signaling to affect gene expression changes. In future work, we plan to explore whether these mechanisms are at play in our system.

A small percentage of the smooth multicellular structures we observe in our P8 and P10 conditions (Fig. 4B) resemble normal lobular and acinar breast structures. Interestingly, Bissell and colleagues previously reported the reversion of a malignant breast cancer cell line into a normal acinar phenotype through the blockade of integrin beta 1 (ITGB1) ^38^. Thus, a link may exist between the short fibril architecture, cell roundedness, and a reduction in the ability of ITGB1 to engage with the matrix. Further, the heterogeneity in the structures formed by the breast cancer cells in our system may represent different integrin-dependent responses as well as different metastatic capabilities. The more abundant network forming phenotype we observe is reminiscent of the collective migration pattern implicated as the primary mode of tumor cell dissemination^39,40^. This collective behavior is thought to be linked to circulating tumor cells that are present as aggregates, which are predictive of poorer clinical outcomes^41,42^. Thus, collagen architecture may influence the metastatic capabilities of cancer cells through modulation of migration phenotype.

Previously, Friedl and colleagues reported that increases in collagen matrix density induced “cellular jamming” in highly aggressive fibrosarcoma and melanoma cells leading to cell chain formation. Their study implicated pore size as the critical matrix feature inducing this migration switching phenomenon^4,43^. The multicellular chain phenotype we observe in our P6 construct is highly similar to that reported by Friedl, where individual cell bodies are distinguishable but connected, resembling a pearl necklace. However, the fused networks, glands, and lobule structures formed in our P8 and P10 conditions are distinct. Further, we find no systematic relationship between pore size and the phenotypic switch. These differences could arise from the fact that we automated the measurement of pore sizes through image processing (see Methods).However, our automated pore size measurement for the 2.5 mg/ml collagen condition (∼2.2 μm^2^) is consistent with that reported previously by other groups using confocal reflection microscopy image analysis (∼0.78-1.8 μm^2^ pore areas for 2.5 mg/ml collagen^44^; 2-5 μm^2^ pore areas for 1.7 mg/ml^45^) and cryo-EM (∼3 μm^2^ pore areas, 2mg/ml collagen^46^). We are also studying a different cell type.

Our successful use of varying concentrations of 8,000 Da PEG to modulate collagen matrix architecture begs the question of whether MC chain size and chemistry could be two additional matrix “tuning knobs” that could be further explored for collagen matrix engineering. The chemical nature of an MC agent as well as its size can determine its exclusion from molecular surfaces. Measurements of the exclusion of different molecular weights of PEG from proteins is largely consistent with the crowding by hard spheres model, where the radii is approximated by the radius of gyration of the PEG polymer^47,48^. However, other measurements have shown that PEG polymers of different sizes are not always excluded from well-defined cavities, as a hard-sphere model would require^49^. A wide range of partial exclusion as a function of molecular size and concentration are possible^50^. This may explain, in part, the reverse in matrix parameter trends we observed in our high concentration, 10 mg/ml PEG, condition. The nonlinearity of the behavior of collagen, a semi-flexible crosslinked biopolymer network, in response to crowding by PEG, is not unexpected. A deep and predictive understanding of such networks has proven to be a daunting theoretical challenge in the field of soft matter and polymer physics. Another example of such non-linear behavior has been demonstrated by crowding actin filaments with PEG, where bundling is induced when the concentration of PEG exceeds a critical onset value^51^. Indeed, the unique characteristics of biopolymers compared to synthetic polymers make their study highly important for fundamental biological understanding.

Also intriguing is our observation that cancer cell proliferation and viability increased slightly when PEG was added after collagen polymerization and maintained in culture for one week. Yet, no significant effect on cell morphology and migration was observed under these control conditions. These finding suggest that molecular crowding may promote proliferation of tumor cells. Previous studies have found that PEG crowding can have the effect of increasing the hydration of proteins and imposing osmotic stress on cells ^48,52,53^. Both increased hydration and osmotic stress have been associated with cancerous tissues ^54,55^. Thus, the addition of PEG to cells in 3D after matrix polymerization could be explored further to probe mechanisms of crowding induced cancer proliferation.

While collagen is only one of many matrix components within the tissue and tumor microenvironment, both clinical and *in vivo* studies have established the relevance of this particular ECM molecule. Collagen is both an independent clinical prognostic indicator of cancer progression ^56^ and a driver of tumorigenesis and metastasis ^57^. As such, understanding how 3D collagen regulates cancer cell migration behavior is likely to provide useful insights into disease pathogenesis. A goal of future work will be to explore in molecular detail the connections between these 3D collagen *in vitro* cancer phenotypes and clinical patient tumors rich in collagen.

## Conclusions

A deeper understanding of the microenvironmental regulators of cancer cell migration could help identify therapies to combat metastasis and improve patient outcomes. By decoupling matrix architecture from matrix density and stiffness, our study identifies a novel role for collagen architecture in modulating cancer cell behavior. The techniques developed herein to modulate collagen architecture allowed us to identify relationships between fibril length, cell shape, and migration phenotype. The same techniques could be extendable to investigations of metastatic migration *in vivo,* since our 3D matrix constructs are collagen and PEG-based, non-toxic, and implantable. These techniques may also be useful for stem cell and regenerative medicine studies as a means to control 3D cell shape and morphogenesis outcomes.

## Methods

### Cancer cell culture

MDA-MB-231 breast cancer cells were ordered from ATCC (Manassas, VA) and cultured in Dulbecco’s Modified Eagle’s Medium (Life Technologies, Carlsbad, CA) supplemented with fetal bovine serum (Corning, Corning, NY) and gentamicin (Life Technologies, Carlsbad, CA), at 37°C and 5% CO_2_. Culture media was changed every other day as needed. Cells were cultured to confluence prior to being trypsinized and embedded inside of 3D collagen I matrices. Cell laden gels were cultured for a week to observe long-term phenotypic differences.

### 3D Culture in Collagen I Matrix

High concentration, rat tail acid extracted type I collagen was ordered form Corning (Corning, NY). MMC agents, PEG 8000 (8,000 Da) and Ficoll 400 (400,000 Da), were ordered in powder form from Sigma-Aldrich (St. Louis, MO) and reconstituted in PBS (Life Technologies, Carlsbad, CA) prior to usage. Trypsinized cells to be embedded, were first mixed with 1x reconstitution buffer composed of sodium bicarbonate, HEPES free acid, and nanopure water. Appropriate amounts of MMC agent, PEG or Ficoll, were then added to produce final concentrations of 0, 2, 4, 6, 8, and 10 mg ml^-1^ PEG (denoted by P0, P2, P4, P6, P8, P10) or 25 mg ml^-1^ Ficoll (denoted by F25). Afterwards, collagen solution was added to the mixture for a final concentration of 2.5 mg ml^-1^ collagen. Finally, pH of the final mixture was adjusted using 1N sodium hydroxide, prior to polymerization via incubation at 37°C (∼20-30 minutes). Gels were prepared inside of 48-well plates with a total volume of 200μl. Following gel polymerization and solidification, MMC molecules were washed out of the collagen gels by rinsing with PBS 3x for 5 min each. Cell culture media was then added on top of the gels after and changed every two days as necessary.

### MMC control experiments

To ensure that the MMC molecules being used were truly inert, we investigated any potential effect that the MMC molecules may have on cells independent of the changing matrix architecture. We conducted a series of MMC control experiments, where MDA-MB-231 cells were embedded inside of three 2.5 mg ml^-1^ collagen gels. After the gels had polymerized, normal culture media was added on top to one of the gel, while the other two either had 10 mg ml^-1^ PEG or 25 mg ml^-1^ Ficoll added into the media on top. The MMC molecules were left in the media and allowed to diffuse down into the gel with the cells. Subsequent media changes would also include appropriate amounts of MMC agent to maintain the PEG and Ficoll concentrations in the gel.

### Confocal reflection imaging of collagen architecture

Collagen matrix architecture and topography was investigated by imaging gels using confocal reflection microscopy (CRM) using a Leica SP5 inverted confocal microscope, equipped with a 20X immersion objective (NA=1.0). Collagen fibrils were imaged by exciting with and collecting backscattered light at 488nm. Confocal reflection imaging is restricted to fibrils that are oriented within 50^°^ of the imaging plane^58^. To verify that the gels are isotropic in their fibril structure, we imaged a gel from the top and from one of the sides (XY and YZ planes), using the same imaging settings. Supplementary Figure 4, A-C shows that differences were negligible.

### Time lapse imaging microscopy

Time lapse microscopy was conducted using a Nikon Ti-E inverted microscope, equipped with a stage top incubation system, to analyze cell motility and migration behavior, morphology, and proliferation and viability. Cells were allowed to settle in the collagen gel in the incubator for approximately 7 h after gel polymerization; time-lapse imaging began at around the 8^th^ hour after the cells were embedded into the collagen gels. Each gel was imaged over 6 fields of view (FOV) for a period of 15 h, with images being taken every 2 min.

### Cell proliferation assay

Cell viability was assessed using a Live and Dead Cell Assay (Abcam, Cambridge, UK). Intact, viable cells fluoresce green (imaged under FITC channel) while dead cells fluoresce red (imaged under TRITC channel). The average number of live cells, dead cells, and live cell viability percentages were calculated over 4 FOVs per condition. Live cell viability % is defined as the number of live cells / the total number of cells * 100.

### Matrix analysis

All matrix analyses were done using the CRM images (fiber length and pore size) and SEM images (fiber width) of the collagen gels in each condition over 3 FOVs. Fibril analysis was conducted in CT-FIRE v1.3 by measuring individual fibril length and width as previously published^59^. Minimum fibril length, dangler length threshold (thresh_dang_L), short fibril length threshold (thresh_short_L), distance for linking same-oriented fibrils (thresh_linkd), and minimum length of a free fibril (thresh_flen), were all set to three pixels. Default settings were used for all other fibril extraction parameters and output figure controls. Settings were optimized to detect and analyze discrete fibrils in CRM images on scales of 0.72μm per pixel. Examples of the fibrils found by CT-FIRE are shown in Supplementary Figure 4D, along with the associated reflection confocal micrographs.

Pore size was calculated using NIS-Elements software (Nikon) as the 2D area encompassed by fibrils. The total pore area (ε’) as a 2D approximation of the 3D tissue ultrastructure (ε) can be described as:

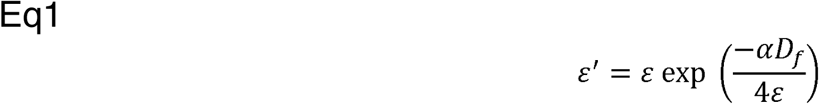

as long as the stereological assumption is met^60^. This assumption was validated by imaging XY and YZ planes of a 2.5 mg/ml collagen gel and analyzing the fibril length (no significant difference), pore area (<0.5μm^2^ difference), which are shown in Supplementary Figure 4, A-C. Additionally, if the imaging depth of field (D_f_) is small enough ε′ = ε.We calculate our D_f_ from:

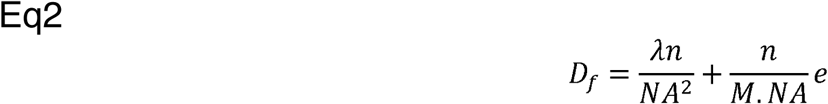

to be ∼0.67 microns. This is smaller than the pixel size in our system and smaller than the expected ε, which makes the fraction in Eq 1 < 1 and suggests that ε′ ≌ ε is a valid approximation. These analyses suggest that our 2D confocal micrographs are a close representation of the 3D architecture. Pre-processing of images were conducted by implementing a Gauss-Laplace Sharpen set to a power of 2 and then by using a Rolling Ball Correction with a rolling ball radius of 15^61^. The contrast of all images were equalized, then images were binarized by thresholding to the same range. Next the automated measurement tool was used to measure pore areas in the binarized images. Single pixel pore values were attributed to speckle noise and removed from all conditions. Homo-/heterogeneity of the matrices was characterized by calculating the coefficient of variation (CV) of fibril length and pore size data distributions.

### Rheometry

Rheological measurements were conducted following established protocols ^27,62^ using a TA Instruments AR-G2 Rheometer. A parallel-plate geometry (20 mm diameter) with a 1000μm gap height was used on 500 μl collagen/PEG gels. For each condition, a strain sweep at frequency of 1 rad/s was recorded to determine each respective linear viscoelastic region. A 1% strain was determined to fall within linear viscoelastic region of each matrix condition. This strain was then held constant across conditions during subsequent frequency sweep experiments. The storage modulus (G’) and loss modulus (G’’) were recorded over frequencies of 0.25-100 rad/s for each condition. The storage modulus at 1 rad/s was reported for each condition.

*Atomic Force Microscopy (AFM)*

AFM was performed to measure local collagen gel stiffness as previously described^63-66^. Briefly, nano-indentations were performed using a MFP-3D Bio Atomic Force Microscope (Oxford Instruments) mounted on a Ti-U fluorescent inverted microscope (Nikon Instruments). A pyrex-nitride probe with a pyramid tip (nominal spring constants of 0.08 N/m, 35° half-angle opening, and tip radius of 10 nm, NanoAndMore USA Corporation, cat # PNP-TR) was first calibrated for the deflection inverse optical lever sensitivity (Defl InvOLS) by indentation in PBS on glass followed by using a thermal noise method provided by the Igor 6.34A software (WaveMetrics) as previously described^63^. Samples were loaded on the AFM, submersed in phosphate buffered saline (PBS), and indented at a velocity of 2 μm/s. Samples were indented until the trigger point, 2 nN, was achieved. Five measurements, equally spaced 50 μm apart, were taken per gel. Tip deflections were converted to indentation force for all samples using their respective tip spring constants and Hooke’s Law. Elastic modulus was calculated based on a Hertz-based fit using a built-in code written in the Igor 6.34A software. The Hertz model used for a pyramidal tip is

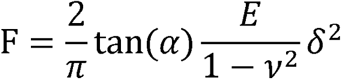

where F is force, α is the half-angle opening of the tip, a is the Poisson’s ratio, E is elastic modulus and δ is indentation. We assumed a Poisson’s ratio of 0.5 (incompressible) for all samples. Curves were only analyzed if at least 80% of the fit curve crossed over the data points. Supplementary Figure 5A shows force-position plots of raw data where the calculated indentation points are indicated and aligned between soft (blue) and stiff curves (red). When the indentation force trigger was met, e.g. when the probe is centered over a pore, data is not collected. Supplementary Figure 5B plots an example of the force curve (orange) that would be obtained from a pore, noting that there is no calculable contact point and tip deflection is only the result of drag on the cantilever during tip approach.

### Scanning Electron Microscope (SEM)

Collagen gels were prepared at 2.5 mg/mL concentration with and without the addition of 10 mg/mL 8KDa PEG, then placed in a humidified incubator (37^°^C) until fully polymerized as described above. The samples polymerized in the presence of PEG were separated into washed and not washed preparations. To wash the PEG after polymerization, PBS was added on top of the gel and placed in the incubator for 5 minutes 3 times. Next, all samples were fixed with 4% PFA for 1 hour at room temperature and the washed 3X with PBS. The samples were then dehydrated by treating them with increasing concentrations of ethanol (50% to 100%). Samples immersed in 100% ethanol were subjected to critical point drying (Autosamdri-815, Tousimis, Rockville, MD, USA), coated with a thin layer of Iridium (Emitech K575X, Quorum technologies, Ashford, UK) and imaged using a Zeiss sigma 500 SEM. Gels were cut with a razor blade to expose cross-sections of the material for imaging.

### Fluorescent PEG analysis

To analyze the penetration of the PEG crowding agent into the 3D collagen matrices, we added fluorescently labelled rhodamine-PEG (10kDa, Nanocs, Boston, MA) to the top of the matrices and conducted fluorescence intensity analysis at the bottom of the matrices over time. For these experiments, 2.5mg/mL collagen gels were prepared in triplicate as described above. After an initial polymerization stage in a cell culture incubator (37°C, 5% CO_2_), the gels were transferred to a bottom reading microplate reader (Tecan, Infinite M1000 Pro) and maintained at 37oC. Baseline fluorescence measurements were performed with excitation at 540nm and emission at 610 nm, corresponding to the spectra of Rhodamine B. After baseline intensities were recorded, culture medium containing 10mg/mL fluorescently labelled PEG was added on top of the gels and fluorescence intensity was recorded at the bottom of the gel by the plate reader every 1 minute for 35 minutes. After this, the PEG solution was removed from the top of the matrices, they were washed three times with PBS, and new fluorescence measurements were performed.

### Cell analysis

Individual cells in the time lapse videos were tracked in Metamorph for motility characterization. Within the 15 h time lapse window, cells were analyzed in terms of the total path length traveled, their average speed, the invasion distance (displacement), and the persistence of their migration (defined as the invasion distance / the total path length traveled). Cell morphology analysis was conducted using images of the cells during the 11^th^ and 15^th^ hour after seeding in the gel, in terms of circularity:

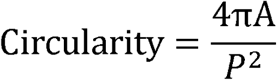

where A is the cross-sectional area and P is cross-sectional perimeter of the cell.

### Correlation analysis

Correlations were calculated in terms of the Pearson correlation coefficient. Correlations were drawn between various matrix parameters to evaluate whether they had been decoupled, as well as between matrix and cell parameters to investigate cell-matrix interactions.

### Gene expression analysis

Biological triplicates of total RNA were prepared for sequencing using the TruSeq Stranded mRNA Sample Prep Kit (Illumina, San Diego, CA) and sequenced on the Illumina MiSeq platform at a depth of >25 million reads per sample. The read aligner Bowtie2 was used to build an index of the reference human genome hg19 UCSC and transcriptome. Paired end reads were aligned to this index using Bowtie2^67^ and streamed to eXpress^68^ for transcript abundance quantification using command line “bowtie2 -a -p 10 -x/hg19 -1 reads_R1.fastq -2 reads_R2.fastq | express transcripts_hg19.fasta”. For downstream analysis TPM was used as a measure of gene expression. A gene was considered detected if it had mean TPM >5.

### Statistics

Data presented in bar graph format was analyzed using one-way analysis of variance (ANOVA) followed by Tukey or Newman-Keuls post hoc test in GraphPad Prism (v5). Coefficient of Variation was calculated by dividing the standard deviation by the mean. Correlation plots were analyzed by Pearson Correlation in GraphPad Prism (v5).

Pearson r correlation coefficient and two-tailed p values are reported. N=3 biological replicates for each condition tested. Statistical significance was set at values of p < 0.05 and reported as p<0.001, ***; p<0.01, **; p<0.05, *.

## Author Contributions

A.H., S.R., and S.F. designed the study. S.R. conducted rheometry experiments. S.R. and A.K. conducted AFM experiments. D.O.V. conducted SEM and fluorescent PEG experiments. D.O.V. and S.R. conducted all other supplementary data experiments.

A.H. conducted all other experiments. All authors contributed to writing or editing the manuscript.

### Acknowledgements

We thank Dr. Karen Christman for use of her rheometer and Jessica Ungerleider for rheometry experimental support. We thank members of the Fraley lab for their technical assistance and insightful comments regarding this work. We thank NSF REU student Tyler Goshia for his assistance in preliminary studies of collagen crowding. The authors declare no competing interests.

S.F. and lab is supported by a Burroughs Wellcome Fund Career Award at the Scientific Interface (1012027), NSF CAREER Award (1651855), ACS Institutional Research Grant (15-172-45-IRG) provided through the Moores Cancer Center, UCSD, and UCSD CTRI, FISP, CRES, and AIM pilot grants. Support was also provided by NIH grants (R01CA206880 to A.J.E.) and by fellowship support via NSF GRFP and NIH T32AR060712 (to A.K.).

### Data Availability

All sequencing data from this study has been deposited in the National Center for Biotechnology Information Gene Expression Omnibus (GEO) and is accessible through the GEO Series accession number GSE101209. All other relevant data are available within the article and supplementary files, or from the corresponding author upon request.

